# Electrical Stimulated GLUT4 Signaling Attenuates Critical Illness-Associated Muscle Wasting

**DOI:** 10.1101/2021.11.11.468054

**Authors:** Alex B Addinsall, Nicola Cacciani, Anders Backéus, Yvette Hedström, Ganna Shevchenko, Jonas Bergquist, Lars Larsson

**Affiliations:** Department of Physiology and Pharmacology, Karolinska Institutet, Stockholm, Sweden; Department of Clinical Neuroscience, Karolinska Institutet, Stockholm, Sweden; Department of Chemistry – BMC, Analytical Chemistry, Uppsala University, Uppsala, Sweden

**Author notes:** Corresponding Author: Lars Larsson, Basic and Clinical Muscle Biology Group, Deptment of Physiology and Pharmacology, Karolinska Institutet, Solnavägen 9, Solna, 171 65.

**Keywords:** Critical Illness Myopathy, Muscle wasting, Glut4 Signaling, TBC1D4, E3 Ligase

## Abstract

**Background:** Critical illness myopathy (CIM) is a debilitating condition characterized by the preferential loss of the motor protein myosin. CIM is a byproduct of critical care, attributed to impaired recovery, long‐term complications, and mortality. CIM pathophysiology is complex, heterogeneous and remains incompletely understood, however loss of mechanical stimuli contributes to critical illness associated muscle atrophy and weakness. Passive mechanical loading (ML) and electrical stimulation (ES) therapies augment muscle mass and function. While having beneficial outcomes, the mechanistic underpinning of these therapies is less known. Therefore, here we aimed to assess the mechanism by which chronic supramaximal ES ameliorates CIM in a unique experimental rat model of critical care.

**Methods:** Rats were subjected to 8 days critical care conditions entailing deep sedation, controlled mechanical ventilation, and immobilization with and without direct soleus ES. Muscle size and function were assessed at the single cell level. RNAseq and Western blotting were employed to understand the mechanisms driving ES muscle outcomes in CIM.

**Results:** Following 8 days of controlled mechanical ventilation and immobilization, soleus muscle mass, Myosin:Actin ratio and single muscle fiber maximum force normalized to cross-sectional area (specific force) were reduced by 40-50% (p< 0.0001). ES significantly reduced the loss of soleus muscle fiber cross-sectional area (CSA) and Myosin:Actin ratio by approximately 30% (p< 0.05) yet failed to effect specific force. RNAseq pathway analysis revealed downregulation of insulin signaling in the soleus muscle following critical care and GLUT4 trafficking was reduced by 55% leading to an 85% reduction of muscle glycogen content (p< 0.01). ES promoted phosphofructokinase and insulin signaling pathways to control levels (p< 0.05), consistent with the maintenance of GLUT4 translocation and glycogen levels. AMPK, but not AKT, signaling pathway was stimulated following ES, where the downstream target TBC1D4 increased 3 logFC (p= 0.029) and AMPK-specific P-TBC1D4 levels were increased approximately 2-fold (p= 0.06). Reduction of muscle protein degradation rather than protein synthesis promoted soleus CSA, as ES reduced E3 ubiquitin proteins, Atrogin-1 (p= 0.006) and MuRF1 (p= 0.08) by approximately 50%, downstream of AMPK-FoxO3.

**Conclusions:** ES maintained GLUT4 translocation through increased AMPK-TBC1D4 signaling leading to improved muscle glucose homeostasis. Soleus CSA and myosin content was promoted through reduced protein degradation via AMPK-FoxO3 E3 ligases, Atrogin-1 and MuRF1. These results demonstrate chronic supramaximal ES reduces critical care associated muscle wasting, preserved glucose signaling and reduced muscle protein degradation in CIM.

## Introduction

CIM is a debilitating condition acquired in critical care, characterized by preferential myosin loss, severe muscle wasting and decreased muscle membrane excitability [1]. Approximately 30% of the general critical care population develop CIM, however the presence of sepsis, systemic inflammation and multiorgan dysfunction significantly increase the incidence of CIM to 70-100% [2, 3]. CIM delays recovery and increases morbidity rates, impairs quality of life for survivors and provides a severe drain on the healthcare system [4, 5]. The long-lasting impact of CIM may be currently underappreciated, as controlled mechanical ventilation (CMV) is vital to patient survival during the COVID-19 global pandemic. In our laboratory 15/16 COVID-19 patients experiencing bed rest and CMV have significant pathological preferential myosin loss, the hallmark of CIM (Unpublished results, Lars Larsson).

CIM pathophysiology is complex, heterogeneous among patients and remains incompletely understood. To date, cellular and inflammatory stress imbalance, mitochondrial dysfunction, protein degradation, impaired EC-coupling, insulin resistance and hyperglycemia are implicated in the development of CIM, demonstrating the diverse systems effected during critical care [6-9]. This may be explained in part as critical care encompasses CMV and immobilization in critically ill patients of diverse clinical backgrounds and pharmacological interventions.

The loss of mechanical stimuli, termed mechanical silencing, is a significant driver of muscle atrophy and associated weakness [10]. Early mobilization counters mechanical silencing, relieving the negative effects of critical care [10, 11]. However, many critically ill patients cannot engage in physical therapy due to illness severity, delirium, sedation, and neuromuscular blockade. Our laboratory and others have demonstrated the positive effects of passive mechanical loading (ML) on muscle mass and function in experimental [10, 11] and clinical studies [12, 13]. Specifically, ML augments muscle CSA and specific force of fast and slow twitch muscle in our experimental rat ICU model, while patients subject to ML increased specific force, but not size of slow twitch tibialis anterior following 9 days intervention [10-12]. These benefits may result from contractile protein retention, reduced oxidative stress or post translational modification of myosin.

Electrical stimulation (ES) promotes muscle mass retention and function in patients with CHF [14], COPD [15] and the critically ill [16]. ES simulates exercise activity, promotes muscle growth and strength, and modulate myopathies [17, 18]. In critically ill, ES drives retention of fast twitch muscle fiber size [9] and muscle strength, resulting in reduced weaning time [16]. While others have shown little effect on muscle volume, systematic review reveals the benefit of ES for muscle mass and strength overall [19].

ES therapy has been widely used in different clinical populations, however the mechanisms driving these benefits are less known. Muscle mass and function are typically measured in indirect clinical measures which lack sensitivity. Further mechanistic studies are required to ascertain how these therapies promote muscle mass and function in CIM. The aim of this study is to 1) examine the effect of chronic direct supramaximal ES in our experimental ICU model, where rats are exposed to CMV and post-synaptic neuromuscular blockade and 2) ascertain the mechanisms driving ES associated benefits. Here, we hypothesize that ES will promote skeletal muscle mass and function retention during 8 days of critical care.

## Methods

### Animal ICU Model

All aspects of this study were approved by Karolinska Institutet ethical committee (N263/14) and conducted in accordance with their ethical standards for animal research. Adult female Sprague Dawley rats were divided into control (CNT; n=8) or experimental groups exposed to deep sedation with isoflurane, pharmacologically paralyzed post-synaptically with alpha cobra toxin, and CMV as previously described [7], for 8 days (8D; n=7) or 8D with supramaximal soleus muscle ES (8DES; n=10). While deeply sedated, hindlimbs of 8DES rats underwent surgical implantation of two Teflon-coated multistranded steel wires (Figure 1A). The wires were placed across the soleus muscle, anterior and posterior of the motor point [20]. Wires were connected to a constant current stimulator providing supramaximal bipolar 0.2ms stimuli at 10-30mA. Activation threshold was monitored throughout. Stimulation was performed for 10s, every 20s at 20Hz for 12h per day. One limb of the 8DES animals was stimulated (STIM), while the contralateral limb was a sham-operated control (SHAM). Deeply sedated rats were weighed and euthanized by heart removal. Directly after soleus muscle was excised collected for downstream analysis, followed by all other tissues of interest. Detailed description of our experimental rodent model can be found in supplement.

**Figure 1.**
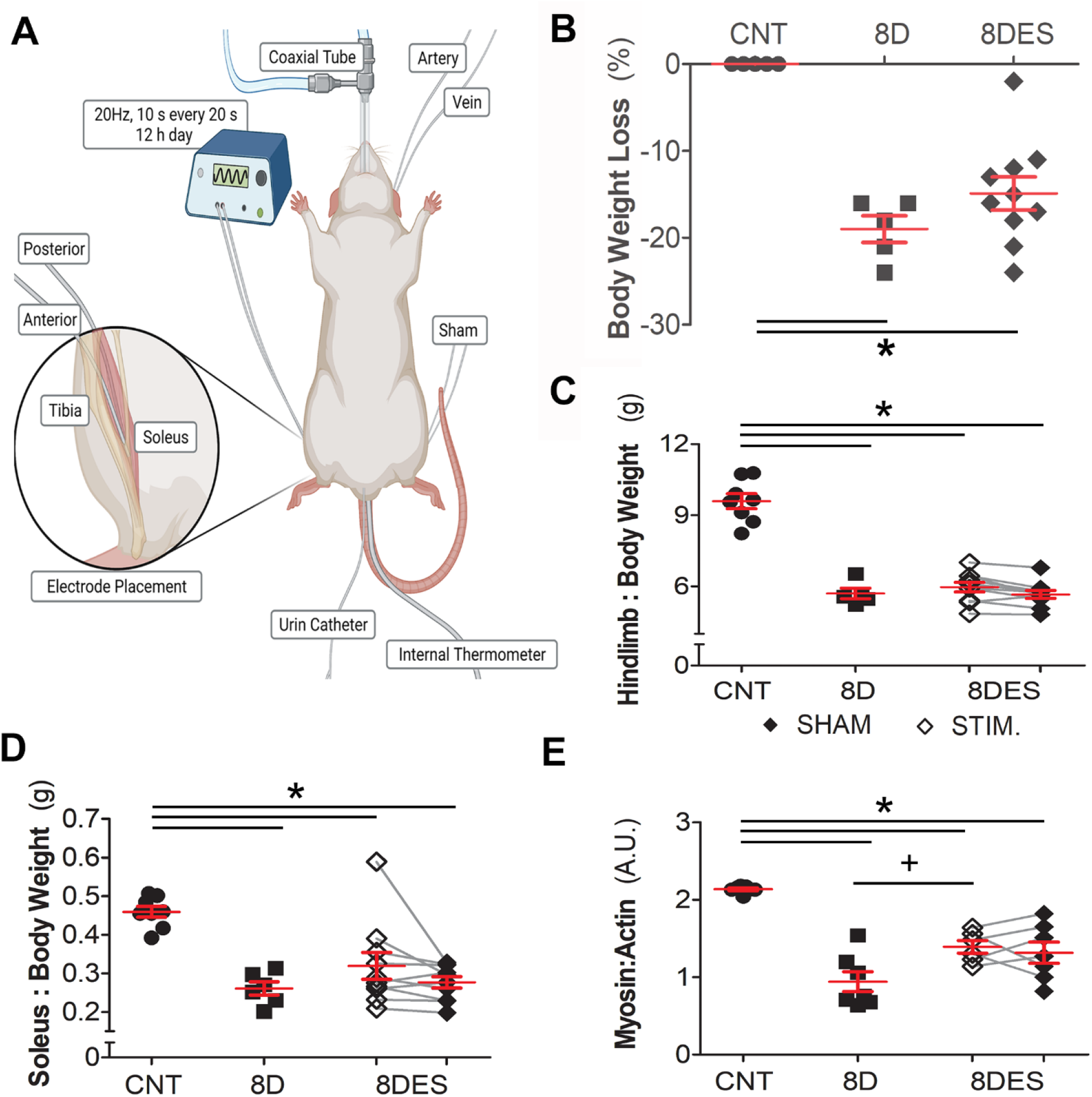
Chronic supramaximal ES promoted Myosin content retention during ventilation. Adult female Sprague Dawley rats were placed in critical care environment entailing deep sedation, CMV, and immobilization for 8 days with (8DES; Stimulated (STIM; open diamonds), Sham control (SHAM; closed diamonds); n = 9) and without (8D; closed squares; n =7) supramaximal ES and compared with control (CNT; filled circles; n = 8). A) Diagram of the experimental model and electrode placement used in this study (Insert). B) Percent body weight loss. C) Hindlimb weight relative to body weight. D) Soleus muscle weight relative to body weight. E) Ratio of Myosin:Actin protein densitometry (n = 5-7 per group). *p<0.05 significantly different from control and +p< 0.05 significantly different with 8DES, as determined by one-way ANOVA.

### Contractile Measurements

Soleus muscles were excised, weighed, and the mid-belly region placed in relaxing solution for muscle bundle preparation [6]. Remaining muscle was snap frozen and stored at −140°C for downstream analysis. Bundles were membrane-permeabilized and cryoprotected before freezing. Cryoprotectant was removed before measuring contractility, where a single muscle fiber segment was isolated and attached between connectors in the setup. Fiber CSA, maximum force, and specific force (normalized to CSA) were calculated as the difference between maximal isometric force and resting tension. A detailed description of the single fiber preparation and contractile measurements is found in supplement.

#### Myosin:Actin Ratio

Myosin and Actin contents were separated on SDS-PAGE, stained with SimplyBlue SafeStain and densitometry quantified as previously described [6]. Results are presented as M:A Ratio.

### RNASeq

RNA was extracted from the soleus as previously described ([6] and supplementary methods), and strand-specific sequencing libraries prepared using NEBNext Ultra RNA Library Prep. Libraries were sequenced (Illumina platform; HiSeq Xten), mapped to the rattus *norvigicus* Ensemble reference genome using HISAT2 [21] and analyzed using EdgeR and Goana pipeline on R [22]. Genes with a false discovery rate (FDR) of p≤ 0.05 were considered significantly differentially expressed genes (DEGs).

### Immunofluorescence

Soleus cross sections were cut, fixed in Acetone, permeabilized and blocked with 1% BSA before co-reacted with anti-GLUT4 and anti-dystrophin primary antibody (Supplementary Table 1). After which, samples were exposed to isotype specific anti-rabbit or mouse Alex Fluora fluorescent secondary antibodies and nuclei counterstained with Dapi. No primary antibody and no secondary antibody controls were included to confirm staining specificity. Representative images were captured by Zeiss LSM800 microscope at ×100 magnification. GLUT4 translocation was quantified by the colocalization of the green fluorescently labelled GLUT4 with the red fluorescently labelled dystrophin, a recognized plasma membrane protein and marker. Image pro software (Media cybernetics; Version 10.0.4) was used to assess the ratio of GLUT4 colocalization area relative to the red fluorescent membrane staining area (µm). All staining and imaging were performed in one batch, to standardised imaging parameters and analysed with a standardised threshold for green and red fluorescence. An average was generated per sample from 3 images at 100x magnification. n=3-4 per group.

### Periodic acid–Schiff (PAS) Immunohistochemistry

Soleus cross sections were cut, fixed in Carnoy’s fixative, and incubated in 0.5% Periodic Acid (Sigma). After which sections were exposed to Shiff’s reagent (Sigma) and nuclei counterstained with Mayer’s hematoxylin (Sigma). Sections were then dehydrated and mounted. Representative images were captured by light microscopy at ×400 magnification.

### Glycogen content

10 mg soleus tissue was homogenized in ddH2O, and supernatant collected before glycogen content was measured using Glycogen Assay Kit (Abcam; ab65620) according to manufacturer’s instructions. Glycogen content was normalized to total protein content (µg/µl) following BCA assay (Pierce, Sweden).

### Western Blotting

Soleus muscle was homogenized in T-Per lysis buffer (ThermoFisher Scientific, Sweden) with phospho, and protease inhibitors (Roche, Sweden) and protein content assessed by BCA assay (Pierce) before 15 ug protein lysates were separated on a TGX Stain-Free™ criterion gel (Bio-Rad, Sweden). Following which, the gel was activated, and proteins visualized using the Chemidoc™ XRS system (Bio-Rad). Proteins were then transferred to PVDF membranes and blocked with 5% BSA or 5% non-fat milk in TBS containing 0.1% Tween-20, before being incubated in primary antibody (Supplementary Table 1) overnight at 4°C. The next day, membranes were washed and subjected to HRP-linked secondary antibodies incubation. Blots were imaged using ECL chemiluminescence (ThermoFisher Scientific). Band densitometry was performed on the Western blots and on the respective stain-free gels using Image lab software (Bio-Rad). Protein expression was normalized to total protein expression.

### Statistics

Statistical analysis was performed using the GraphPad Prism software. One-way ANOVA with Tukey’s post hoc test was used to compare treatment groups. p<0.05 was considered statistically significant. Data are presented as mean ± SEM.

## Results

### Chronic supramaximal ES maintains Myosin content

Loss of body and muscle mass, preferential myosin loss, and force deficit are observed in CIM [23]. After 8D body weight was reduced by 19% (Figure 1B; p< 0.0001) and remained reduced by 13% when ES was administered (p< 0.0001). As a proxy measure of lean mass, gastrocnemius, plantaris, TA, extensor digitorum longus, and soleus hindlimb muscle weights were combined. Hindlimb weight relative to body weight was reduced by 40 and 37% following 8D and 8DES, respectively (Figure 1C; p< 0.0001). Soleus muscle mass was significantly reduced following 8D (38%; Figure 1D; p< 0.0001). ES had no effect on soleus muscle weight. Myosin:Actin ratio is a key clinical feature of CIM [1]. Soleus Myosin:Actin ratio was reduced by 56% during 8D (Figure 1E; p< 0.0001). ES reduced the loss of Myosin:Actin ratio by 33% from 8D (p< 0.05). Myosin:Actin ratio did not differ between stimulated and unstimulated contralateral sham control after 8DES.

### Chronic supramaximal ES lessens soleus wasting

To ascertain the effect of ES on functional performance, soleus muscle fiber CSA and specific force were determined. A total of 218 soleus muscle fibers passed selection criteria (Supplementary Methods). Soleus muscle fiber CSA was reduced by 42% following 8D (Figure 2A; p< 0.0001). ES improved soleus CSA by 27 and 29% from 8D and contralateral sham control, respectively (Figure 2A; p< 0.05). Soleus muscle fiber specific force was reduced by 50% following 8D yet ES failed to effect it (Figure 2B; p< 0.0001). Thus, ES benefits muscle size, but not force generation in CIM.

**Figure 2.**
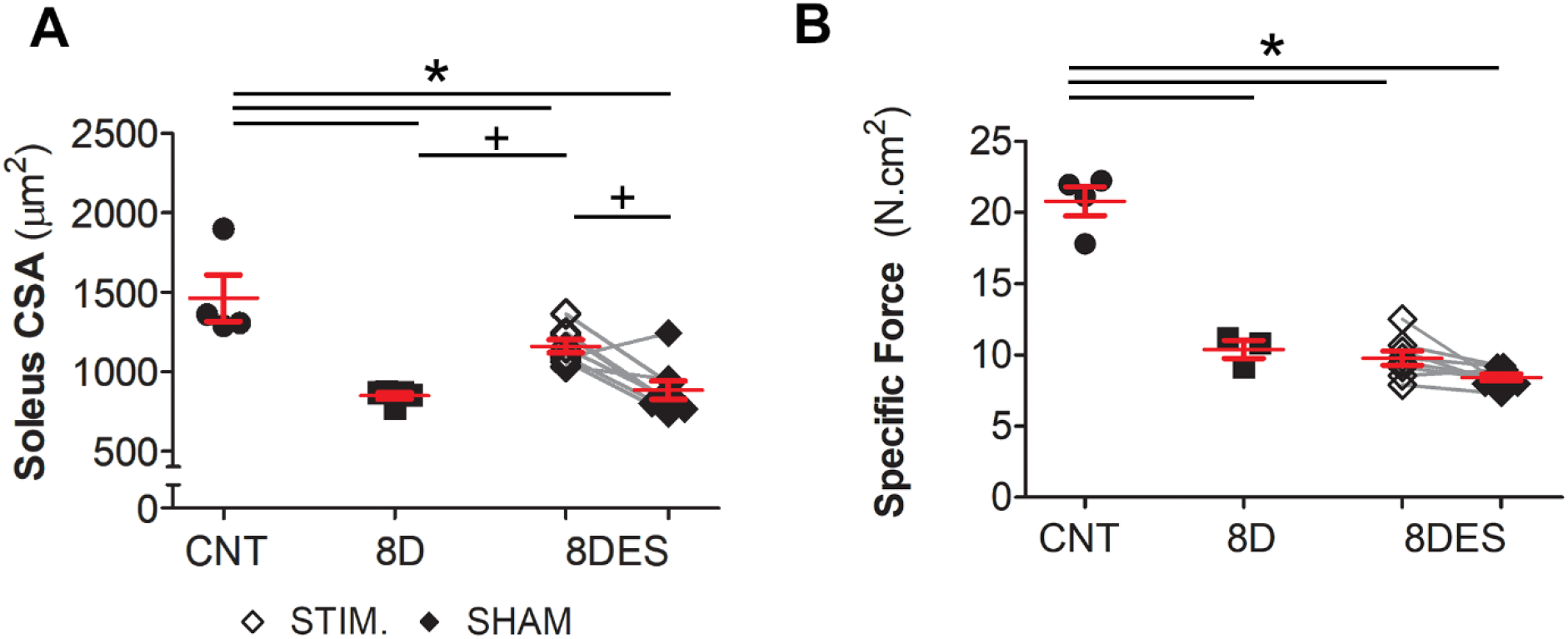
Chronic supramaximal ES ameliorates soleus CSA in CIM. Adult Sprague Dawley rats were placed in critical care environment entailing deep sedation, CMV, and immobilization for 8 days with (8DES; Stimulated (STIM; open diamonds), Sham control (SHAM; closed diamonds); n = 8) and without (8D; closed squares; n =5) supramaximal ES and compared with control (CNT; filled circles; n = 4). A) Soleus muscle fiber cross sectional area (CSA) and B) Soleus muscle fiber force relative to CSA (Specific Force) represented as average per animal. *p< 0.05 significantly different from control or +p< 0.05 significantly different with 8DES, as determined by one-way ANOVA.

### Chronic supramaximal ES preserves glucose signalling

To determine the mechanism by which ES protects soleus fiber size RNASeq was employed. Here, 8D resulted in 2,923 DEGs compared with CNT (p< 0.05; up: 1,340; down: 1,583). The addition of ES resulted in a total of 69 DEGs compared with 8D (fold change ≥ 1.5; p< 0.05), the majority of which were upregulated (61 vs 8; Supplementary data). Enriched Gene Ontology (GO) terms of these upregulated genes were explored. ES enriched GO terms pertaining to *channel activity, action/membrane potential,* and *muscle contraction* (Figure 3A). *Fructose-6-phosphate binding* and *phosphofructokinase activity* were the most enriched terms, indicating ES promoted glycolysis. The DEGs within these terms were muscle specific *Phosphofructokinase 1(PFKM)* and *Phosphofructokinase 2 (Pfkfb1*; Figure 3B). These key glycolysis genes are promoted by increased glucose uptake following exercise.

**Figure 3.**
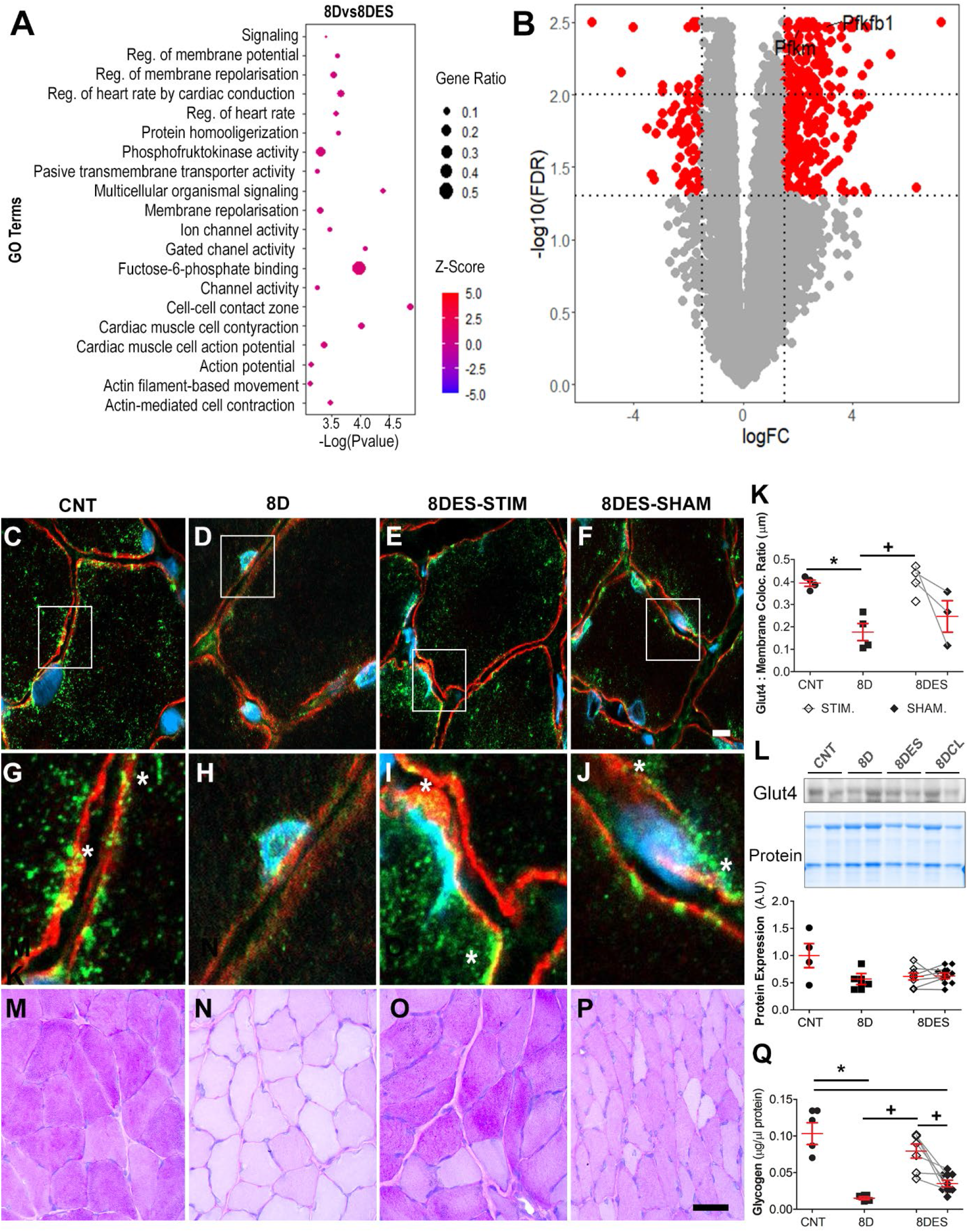
Chronic ES promotes Glut4 translocation. Adult Sprague Dawley rats were placed in critical care environment entailing deep sedation, CMV, and immobilization for 8 days with (8DES; Stimulated, Sham control (n = 4) and without (8D; n =4) supramaximal ES and compared with control (CNT; n = 4). A) Gene ontology (GO) terms were assessed from the upregulated DEGs. B) DEGS from *Phosphofructokinase activity and Frucose-6-phosphate binding* GO terms. C-F) Representative immunofluorescent GLUT4 (green), dystrophin (red) and DAPI (blue) colocalization in soleus muscle cross sections, Scale = 5 µm. G-H) Zoomed sections of the region of focus in C-F (white box). K) Quantification of GLUT4 colocalization relative to sarcolemma area. L) Representative Western blot and graph showing densitometry measured in arbitrary units (A.U) of total GLUT4 protein expression. M-P) Representative PAS histochemical staining, Scale = 50 µm. Q) Muscle glycogen content (n = 4-8). *p< 0.05 significantly different from control or +p< 0.05 significantly different with 8DES, as determined by one-way ANOVA.

In skeletal muscle, GLUT4 is the primary receptor for glucose endocytosis. Immunofluorescent assessment of GLUT4 translocation and quantification of its colocalization at the sarcolemma were performed to examine alteration of glucose uptake (Figure 3C-K). GLUT4 translocation was reduced by 55% following 8D, with GLUT4 staining restricted to the perinuclear region (p = 0.003; Figure 3D, H and K). 8DES preserved GLUT4 translocation, identified by colocalization at the sarcolemma (yellow staining; Figure 3E, I, K). Interestingly, GLUT4 translocation tended to increase in the sham operated contralateral limb yet failed to reach statistical significance (Figure 3D, J, K). Total GLUT4 protein expression remained unchanged following 8D or 8DES (Figure 3L).

To assess the implication of preserved GLUT4 translocation we assessed total glycogen content via PAS stain and glycogen assay. Here, 8D reduced total glycogen levels by 85% (p < 0.001; Figure 3M-Q), The addition of ES increased glycogen content in line with preserved GLUT4 signaling such that glycogen levels were not different from control (Figure 3Q). Glycogen content also tended to increase in the sham operated contralateral limb yet failed to reach statistical significance. Thus, retention of GLUT4 translocation following ES preserves muscle glucose homeostasis.

### AMPK stimulates GLUT4 translocation

Skeletal muscle glucose uptake is primarily driven via insulin stimulated-AKT signaling, contraction induced phosphorylation of AMP-activated protein kinase (AMPK) or increased calcium signaling [24]. To ascertain the mechanism by which GLUT4 translocation is impaired following critical care and preserved with ES, DEGs pertaining to *Insulin signaling* were explored. Genes involved in the insulin stimulated AKT signaling were identified in the top DEGs (Figure 4A; Supplementary Table 2). Here, 8D reduced *Irs1, Inpp5a, Tbc1d4, Flot1, Fbp2* and *Cblb,* while ES promoted these genes (p< 0.05). The activation of this pathway was assessed, yet phosphorylation of AKT at s473 and t308 were not significantly altered following 8D or 8DES (Figure 4B).

**Figure 4.**
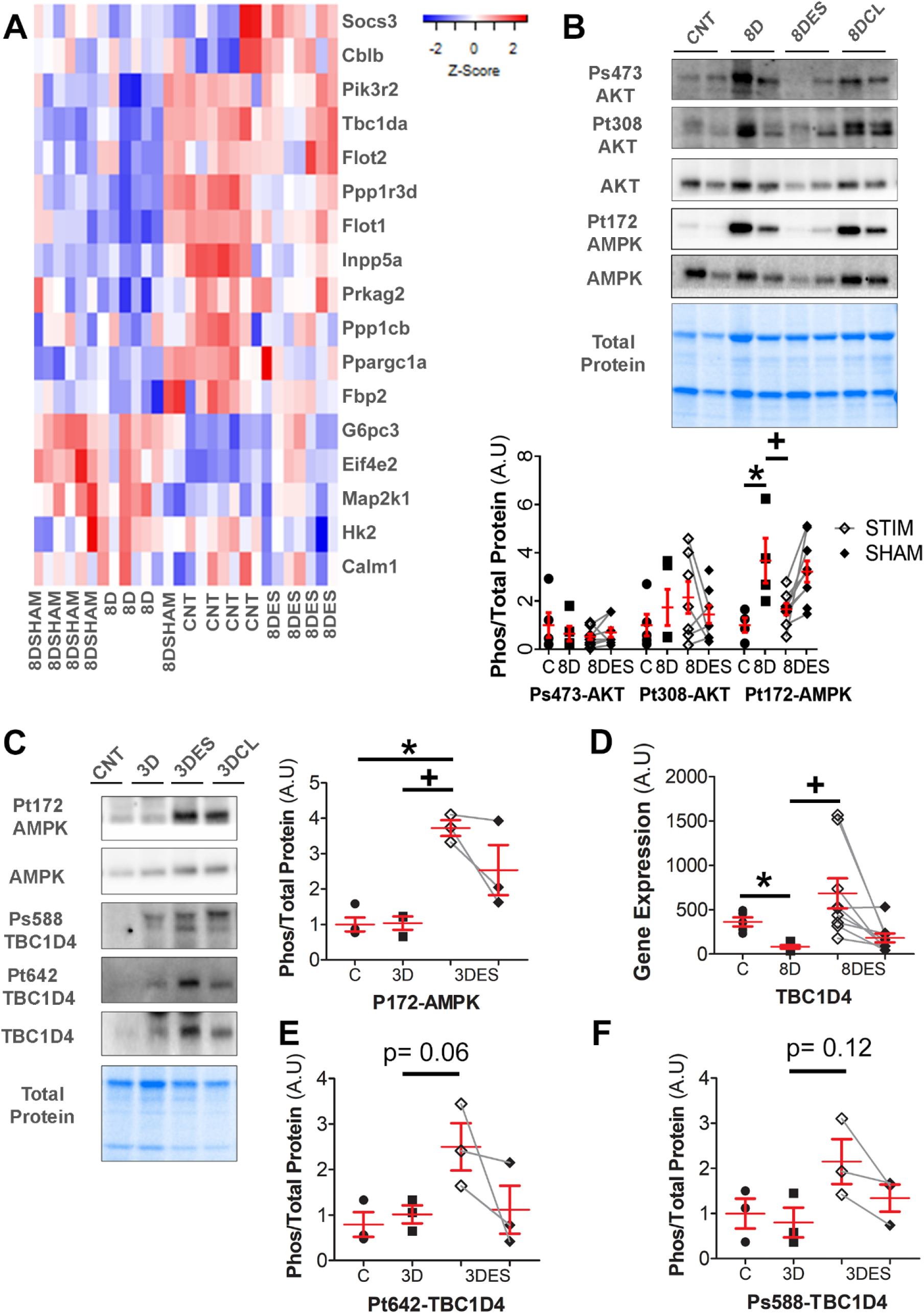
GLUT4 translocation is independent of insulin stimulated -AKT signaling pathway. Adult female Sprague Dawley rats were placed in critical care environment entailing deep sedation, CMV, and immobilization for 8 or 3 days (8DES; n = 9 or 3DES; n = 3; Stimulated (STIM; open diamonds), Sham control (SHAM; closed diamonds)) and without (8D; n = 5 or 3DES; n = 3; closed squares) supramaximal ES and compared with control (CNT; filled circles; n = 5). A) Heat map of top DEGs pertaining to *Insulin Signaling.* B) Representative Western blot with graph showing densitometry measured in arbitrary units (A.U) of P-AKT at s473 and t308, total AKT, P-AMPK at t172, total AMPK and total protein following 8 days ventilation. C) Representative Western blot of P-AMPK at t172, total AMPK, P-TBC1D4 at ser588 TBC1D4 at t642, total TBC1D4 and total protein, with graph showing densitometry measured in A.U of P-AMPK at t172 following 3 days ventilation. D) Transcript expression of TBC1D4 during 8DES. E) Graphs showing densitometry measured in A.U of P-TBC1D4 at t642 following 3 days ventilation. F) Graphs showing densitometry measured in A.U of P-TBC1D4 at ser588 following 3 days ventilation. *p< 0.05 significantly different from control or +p< 0.05 significantly different with 8DES, as determined by one-way ANOVA.

Skeletal muscle contraction can instigate GLUT4 translocation independent of and in addition to insulin stimulated GLUT4 activity, as there are two distinct pools during GLUT4 activation [25]. Skeletal muscle contraction increases calcium release, cytosolic calcium concentration and activates calcium-dependent signaling targets including *calmodulin* and its protein dependent kinases. Indeed, 8DES promotes calcium release and subsequent upregulation of CamK1d and CamKK2 (p= 0.047 and 0.045, respectively; Supplementary Table 3). CaMKK is also known to activate AMPK. Exercise also increases energy utilization, promoting an increased AMP:ADP environment, known to stimulate AMPK. Both increased calcium signaling, and AMPK phosphorylation stimulate GLUT4 translocation. 8D increased P-AMPK at t172, which was subsequently reduced to CNT levels following ES (p= 0.004; Figure 4B). No change was observed in the sham contralateral limb. Lack of AMPK activation following ES was surprising, as acute exercise stimulates AMPK in line with GLUT4 translocation. However, training studies show GLUT4 signaling can be independent of AMPK [26]. Initially, AMPK activates following exercise and initiates GLUT4 translocation. Yet, following continued training GLUT4 translocation can remain active, while P-AMPK may diminish. To examine a time effect with ES, rats were placed in critical care conditions with or without supramaximal ES for 3 days to examine early activation of AMPK. Indeed, 3DES increased P-AMPK from CNT and 3D (p= 0.015, Figure 4C). Interestingly, we observed a trend for increased P-AMPK in the sham contralateral limb following 3D, however it failed to reach statistical significance.

TBC1D4 (also known as AS160) is a downstream target of AKT insulin signaling, yet it’s also activated by AMPK following exercise or contraction at specific phosphorylation sites including s588 and s711, where it potentiates insulin mediated glucose uptake [27, 28]. TBC1D4 transcript expression is reduced by 3 logFC following 8D (p= 0.001; Figure 4D). ES promoted TBC1D4 transcript expression (p= 0.029) while remaining reduced compared to CNT (p= 0.01). To assess general and AMPK relative TBC1D4 activity, phosphorylation of TBC1D4 at t642 and s588 was assessed, respectively. 3DES tended to increase P-TBC1D4 (t642) and P-TBC1D4 (s588) over total TBC1D4 expression in line with P-AMPK activation yet failed to reach statistical significance (Figure 4E and F; p= 0.06 and 0.12). TBC1D4 may be working downstream of AMPK to promote GLUT4 signaling following ES in CIM.

### Chronic supramaximal ES downregulates muscle protein degradation

AMPK is a stress and energy responsive protein implicated in regulation of protein synthesis via mTOR and degradation regulation through FoxO. AMPK is implicated in skeletal muscle anabolism and catabolism. Provided P-AMPK was reduced following 8DES we sought to establish if the increase of soleus muscle CSA was mediated by reduced inhibition of mTOR. Thus, downstream targets of mTOR, S6 protein kinase (S6PK) and 4EBP1 were assessed. Here, P-S6PK and P-4EBP1 were unchanged by 8D or 8DES (Figure 5A and B).

**Figure 5.**
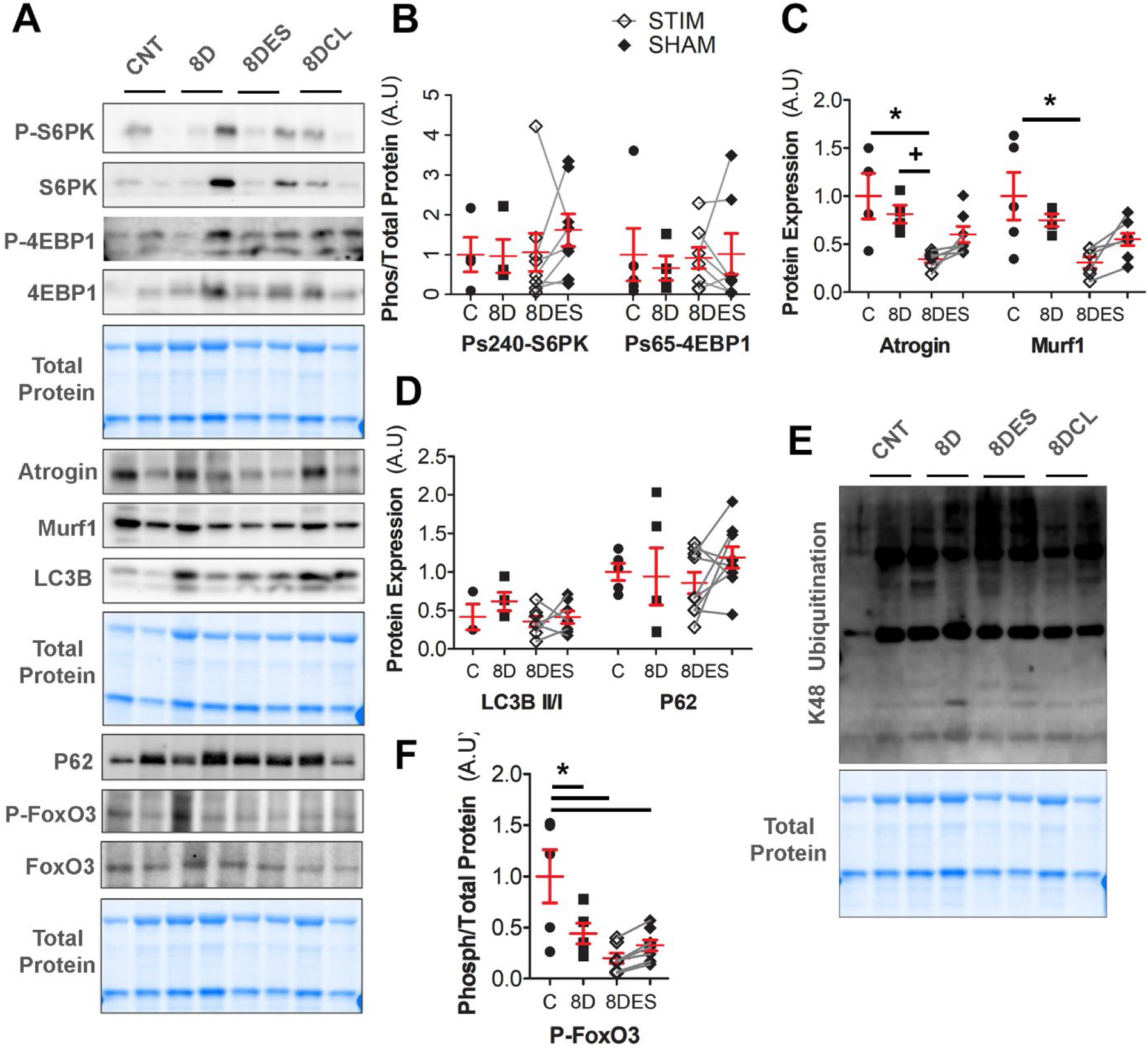
ES reduces muscle E3-ligase activity. Adult female Sprague Dawley rats were placed in critical care environment entailing deep sedation, CMV, and immobilization for 8 days with (8DES; Stimulated (STM; open diamonds), Sham control (SHAM; closed diamonds); n = 9) and without (8D; closed squares; n = 5) supramaximal ES and compared with control (CNT; filled circles; n = 5). A) Representative westerns blotting for key AMPK downstream targets of mTOR and FoxO3. B) Graphs showing densitometry measured in arbitrary units (A.U) of protein synthesis targets P-S6PK and P-4EBP1. C) Graphs showing densitometry measured in A.U of protein ubiquitination targets Atrogin-1 and MuRF1. D) Graphs showing densitometry measured in A.U of protein autophagy targets LC3B and P62. E) Graphs showing densitometry measured in A.U of K48 ubiquitination. F) Graphs showing densitometry measured in A.U of P-FoxO3. *p< 0.05 significantly different from control or +p< 0.05 significantly different with 8DES, as determined by one-way ANOVA.

Downstream AMPK muscle degradative E3-ligases Atrogin-1 and MuRF1, and autophagic proteins LC3B and P62 were assessed. Atrogin-1, but not MuRF1 protein expression was reduced following 8D (p= 0.006; Figure 5A). ES reduced Atrogin-1 and MuRF1 protein levels compared with CNT (p= 0.006 and 0.006, respectively; Figure 5C). 8DES Atrogin-1 and MurRF1 protein levels tended to be reduced from 8D by approximately 50%, yet MuRF1 failed to reach statistical significance (p= 0.08). Interestingly, a non-significant reduction of Atrogin-1 and MuRF1 protein levels were also observed in the sham contralateral limb compared with 8D. The autophagic response was determined by the LC3B II: LC3B I ratio and P62 protein levels. Neither LC3B activation, nor P62 expression were changed following 8D or 8DES (Figure 5D). As Atrogin-1 and MuRF1 can form K48 and 63 ubiquitin linkage for subsequent proteasomal degradation, K48 and total polyubiquitination were assessed. Neither K48 nor total ubiquitination expression were altered by 8D or 8DES, irrespective of the ES associated reduction of Atrogin-1 and MuRF1 (Figure 5E and Supp Figure 2). Nonetheless promotion of soleus CSA and myosin protein content with ES appears associated with reduced E3 ligase activity, but not autophagic response in CIM.

FoxO3 regulates skeletal muscle degradative proteins downstream of AMPK, thus P-FoxO3 activation was examined. Soleus P-FoxO3 was significantly reduced following 8D (p= 0.001; Figure 5F). ES tended to reduce P-FoxO3 compared with 8D yet failed to reach statistical significance.

## Discussion

Critical care is a lifesaving intervention, yet it comes at a cost. CMV, immobility and pharmacological intervention promote the development of morbidities, attributed to impaired recovery, long‐term complications, increased healthcare expense and mortality. The underlying mechanisms are complex and multifactorial. The mechanical silencing unique for modern critical care, defined by a lack of external strain related to weight bearing and internal strain related to activation of contractile proteins is one factor underlying CIM [11]. The deep sedation, delirium, and illness severity limit active engagement in rehabilitation. Targeted physical therapies, passive ML and ES, counter mechanical silencing, and significant promise. However, the mechanisms underpinning these therapies are limited. Here, 8D CIM reduced soleus muscle mass, CSA, specific force, and Myosin:Actin ratio and was associated with reduced GLUT4 signaling and muscle glycogen content. ES augmented soleus muscle fiber atrophy and preferential myosin loss, but not specific force following 8 days critical care. ES promoted GLUT4 signaling and returned muscle glucose homeostasis which corresponded with a trending increase in acute AMPK-TBC1D4 signaling. Improved muscle energy homeostasis was associated with reduced down-stream AMPK-FoxO3 E3-ubiquitin proteins, Atrogin-1 and MuRF1 following 8DES. ES reduced CIM associated reductions in soleus CSA and preferential myosin loss, preserved glucose signaling and reduced muscle protein degradation. These findings are summaries in Figure 6.

**Figure 6.**
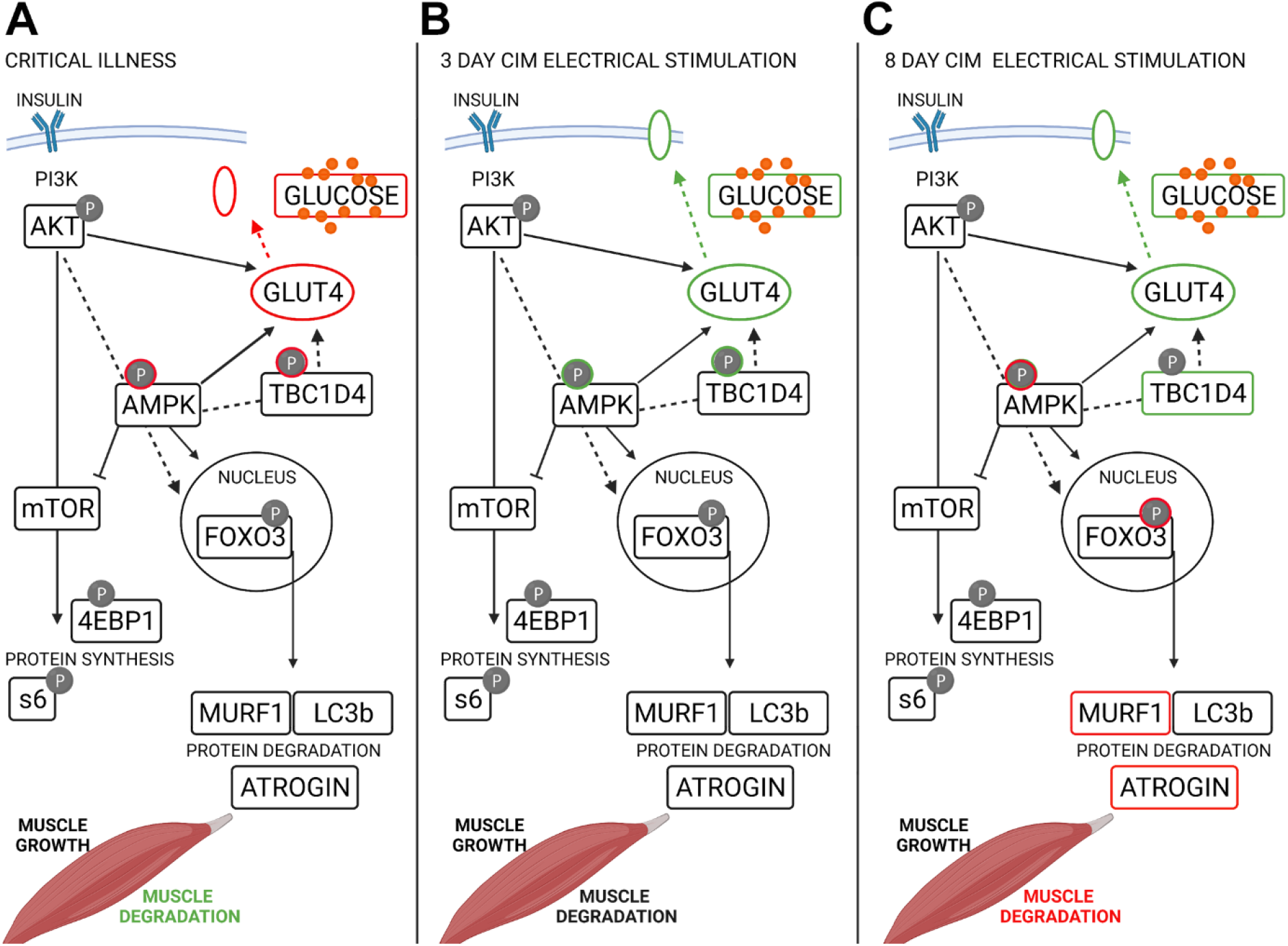
Summary of ES regulation of soleus CSA in CIM. Schematic signaling pathway in A) critical illness, B) 3 day short-term supramaximal ES and C) 8 days long-term supramaximal ES. Red indicates down regulation. Green indicates upregulation.

### Insulin resistance and GLUT4 signaling

Critically ill patients develop insulin resistance and hyperglycemia which contribute to muscle loss and correlate with mortality [29]. Heightened inflammation, reduced incretins, and steroid use contribute, yet impaired GLUT4 signaling was recently attributed to hyperglycemia. Consistent with Weber-Carstens *et al.,* in critically ill patients, we observe impaired GLUT4 signaling in the soleus muscle in our experimental model, where GLUT4 is restricted to the perinuclear space [9]. Impaired GLUT4 translocation could be the result of compromised insulin signaling or lack of mechanical stimuli. Impaired AMPK signaling is attributed to disrupted GLUT4 translocation, as insulin-stimulated AKT activation remained intact in CIM patients [9]. Administration of acute ES for 12 days evoked AMPK stimulated GLUT4 translocation in CIM patients and larger fast, but not slow muscle fiber CSA [9]. The activation of insulin–independent GLUT4 signaling is consistent with our observations in soleus muscle, as P-AKT did not change following ES. The promotions of glucose signaling was enough to overcome the observed deficit of muscle glycogen content with 8D critical care. To the best of our knowledge, this is the first observation of such drastic muscle glycogen reduction with CIM. Thus, AMPK-mediated GLUT4 signaling ameliorates glycogen depletion, returning glucose homeostasis following chronic ES.

AMPK activation can result from increased AMP:ADP ratio or increased cellular stress. Here we measure P-AMPK following acute (3 days) and chronic (8 days) ES. AMPK activity can vary with exercise intensity and duration; however, its downstream processes may remain stimulated [26]. 3DES phosphorylates AMPK, without activation following 3D critical care. However, following 8D we observe the opposite effect, where critical care increases P-AMPK, while ES remained similar to control. One may suggest that these time-related P-AMPK changes shift with the muscle’s energy state. ES AMPK-activated glucose signaling maintains muscle to a homeostatic balance, while 8D remains energy deprived leading to increased P-AMPK in attempts to stimulate glucose uptake or promoting stress-induced AMPK signaling. Reduced P-AMPK following 8DES decreased key skeletal muscle ubiquitination targets Atrogin-1 and MuRF1 downstream of AMPK-FoxO3. While we did not observe any changes in K48 and total ubiquitination downstream of the ES induced reduction of Atrogin-1 and Murf1, it must be acknowledged that hundreds of ubiquitin ligases exist and contribute to ubiquitination beyond these two specific ligases. Future work should examine this relationship in detail. Even so, increased soleus muscle size appears to result in part from reduced E3 ligases, Atrogin-1 and Murf1 following ES.

### The potential of TBC1D4

While ES promotes GLUT4 signaling in CIM patients, whether this stimulates a return of insulin mediated glucose uptake is unknown. Weber-Carstens et al., observed a compensatory increase in AKT phosphorylation in critically ill patients, suggesting insulin-stimulated glucose uptake is impaired downstream of AKT [9]. While we observed no activation of AKT in our model, transcripts of downstream mediator TBC1D4 were significantly reduced following 8D and promoted with ES. TBC1D4 is regulated by AKT during insulin stimulation, mediating skeletal muscle GLUT4 translocation and glucose uptake [30, 31]. TBC1D4 is also activated by AMPK-specific phosphorylation sites following exercise, contraction, or AICAR, where it improves skeletal muscle insulin sensitivity [27, 28]. Promotion of TBC1D4 and AMPK with ES may suggest its involvement in returned GLUT4 signaling in CIM. Whether increased TBC1D4 levels are associated with improved insulin sensitivity and normoglycemia following ES in CIM is unknown and a limitation of this study. Nonetheless, the potential of TBC1D4 in returning homeostatic glucose regulation following muscle contraction in CIM warrants further exploration.

### GLUT4 translocation machinery and CIM

AMPK-TBC1D4 signaling may be disrupted in CIM, however disrupted GLUT4 translocation observed with CIM could also result from impaired exocytosis machinery. RNAseq identified members of the lipid raft responsible for GLUT4 exocytosis and membrane docking to be reduced 8D (Figure 3). The presence of Caveolin-3 and Flotillin-1 are crucial to insulin-stimulated GLUT4 translocation [32]. Caveolin-3-null mice are insulin-resistant and have decreased skeletal muscle glucose uptake [33]. Mutations of Caveolin-3 cause Limb-girdle muscular dystrophy, disrupted sarcolemma and insulin-resistance [34]. Alternatively, disrupted Flotillin-1 prevents GLUT4 translocation, and glucose uptake in response to insulin [32]. Reduction of these lipid raft genes in CIM may contribute to impaired GLUT4 signaling and the development of hyperglycemia in CIM. ES preserved Flotillin-1 expression which coincided with preserved GLUT4 signaling. However, this comes with a caveat, as Flotillin-1 upregulation following ES could result from the exercise-like stimulus, as Flotillin-1 expression is used to identify extracellular vesicle secretion following exercise [35]. ES promoted GLUT4 translocation and muscle glycogen content such that it was no different from control. Interestingly, this occurred irrespective of the loss of Caveolin-3 following ventilation, which was not corrected through ES (Supplementary Figure 1). Caveolin-3 transcript expression is however increased following ML, where its mechanosensory capabilities may contribute to improved muscle outcomes [11]. The implication of ML on GLUT4 signaling and glucose homeostasis is not known in CIM. One may speculate the exercise nature of ML would promote GLUT4 signaling, contributing to the protection of muscle size and function during critical care. Further research is required to investigate the mechanics of the lipid raft and GLUT4 translocation in the context of CIM.

### ES and ML Compared

Passive ML and ES are modalities to maintain muscle mass in patients unable to actively partake in rehabilitating physical therapies. Previous work from our group demonstrates the benefits of passive ML on soleus muscle fiber size, specific force, and preferential myosin loss [11, 12]. ES benefits were less pronounced than ML. This is surprising as ES is expected to show a stronger effect compared with ML, as ES provides more than 30-fold more activations (400/min) then ML (13/min). Additionally, ES activates contractile proteins via intracellular calcium signaling, modulating internal strain and mechanosensitive pathways, while ML primarily stimulates mechanosensation. The force improvement following ML, which was absent with ES, may be facilitated through other mechanisms including regulation of oxidative stress or expression of sarcomeric proteins as these are improved following ML, but not ES [11]. It must also be noted that a high proportion of ES and few sham control muscle fibers presented with fragility upon contractility testing. This fragility is associated with reduced Tuba1b transcript expression following ES, when compared with 8D (−1.97 logFC). Tubulins impart mechanical stability in muscle, are reduced during unloading, yet maintained with passive stretch [36]. The lack of force generation following ES may be explained in part through muscle fragility and this mode of ES delivery may impact its translation to clinic. While these modalities provide muscle performance outcomes by different means, it’s interesting that the combination of active cycling and ES didn’t improve muscle strength in critical care patients [37], suggesting little cumulative effect. While both therapies intend to counter the mechanical silencing associated with critical care, neither of these therapies have restored size and function to normal levels. Nonetheless, results from our groups and others suggest ML to have superior benefit [11, 38].

### Potential systemic effect

Interestingly, the contralateral sham control soleus observed M:A ratio, AMPK, Glut4 signaling and reduction of Atrogin-1 and MuRF1 similar to the ES soleus, albeit to a lesser extent. While many of these failed to reach statistical significance or have a significant effect upon soleus CSA, this cross-over is suggestive of a systemic effect with ES not observed during ML [11]. In emulating exercise, ES promotes the secretory function of skeletal muscle, releasing circulatory factors know as ‘myokines.’ Myokines promote a cascade of endocrine, paracrine, and autocrine effects which modulate inter-organ functions [39]. Thus, myokines exert beneficial effects on a plethora of conditions not limited to muscle growth and insulin sensitizing. One may speculate that the direct ES delivered here could stimulate various myokines or calcium induced small vesicles and exosomes that have beneficial autocrine effect on the contralateral limb of the stimulated rats. The return of Flot1 with ES supports the involvement of ES induced exosomes. Release of circulatory factors can also influence the neuromuscular junction (NMJ). Observations from our group suggest ES also has a strong effect on terminal axon growth and Schwann cell spouting at the NMJ of stimulated and contralateral control limbs (In manuscript, Larsson, Cacciani and Thompson). Why particular elements of this study observe a global effect, while others do not, requires further work and is of interest to conditions like CIM and similar conditions where immobility and sedation limit participation in active physical therapies.

### Conclusion

Our findings demonstrate the benefit of chronic supramaximal ES on soleus muscle glucose homeostasis, size and myosin content during long-term immobilization and mechanical ventilation. The in-depth mechanistic approach identified attenuation of soleus CSA which may in part be regulated by activation of AMPK-TBC1D4 mediated GLUT4 signaling and preserved muscle glycogen content. Maintenance of skeletal muscle glucose homeostasis reduced downstream mediator of E3 ligases, Atrogin-1 and MuRF1 via AMPK-FoxO3 signaling. This study identifies potential targets that may modulate insulin sensitivity in CIM that should be the focus of future research. Overall, this study demonstrates ES as a potential therapy to ameliorate muscle size in the critically ill.

## Ethics Statement

All authors of this manuscript certify that they comply with the Ethical guidelines for authorship and publishing in the Journal of Cachexia, Sarcopenia and Muscle [40].

## Acknowledgement

This study was supported by grants from the Swedish Research Council (8651, 7154, 1001I, the Swedish Heart and Lung Foundation, the Erling-Persson Foundation, Stockholm City Council (Alf 20150423, 20170133), Centrum för Idrottsforskning (2020-0014; 118-2021), and Karolinska Institutet to LL and Centrum För Idrottsforskning Postdoctoral Fellowship (D2020-0018) and Loo and Hans Osterman’s foundation to ABA. We appreciate the kind gift of the AKT, TC1D4 and AMPK antibodies from Dr. Victoria Wyckelsma, Dr. Alexander Chibalin and Ms. Alice Maestri of Karolinska Institutet.

## Conflict of interest

The authors declare no conflict of interest.

## Author contributions

The experimental work was performed in the research laboratory of LL in the Department of Physiology and Pharmacology, Karolinska Institutet. LL and ABA conceived the study design. LL, NC, AB, YH and ABA conducted the animal experiments. ABA, NC, GS and JB undertook down-stream analysis. ABA wrote the manuscript in conjunction with NC and LL. All authors approved the final version of manuscript.

## Notes

### Competing Interest Statement

The authors have declared no competing interest.

### Summary of Updates

Figure 4 updated to clarify AKT phosphorylation.

