## Supplementary file for "Electrical Stimulated GLUT4 Signaling Attenuates Critical Illness-Associated Muscle Wasting" (from another version of this preprint: version 2, 2021-12-06)

**Supplementary Results**

**Supplementary Table 1. Antibodies.**

| **Target** | **Concentration** | **Details** |
|---|---|---|
| *Glut4* | IHC: 1:200; WB: 1:1000 | *2213, Cell Signaling Technology*, USA |
| *Dystrophin* | IHC: 1:50 | *Sc-15376, Santa Cruz Biotechnology, USA* |
| *P-s473 AKT* | 1:1000 | *Ab 81283, Abcam, USA* |
| *P-t308 AKT* | 1:1000 | *4056, Cell Signaling Technology* |
| *AKT* | 1:1000 | *9272, Cell Signaling Technology* |
| *Beta Actin* | 1:1000 | *Ab8227, Abcam* |
| *P-t172 AMPK* | 1:1000 | *2535, Cell Signaling Technology* |
| *AMPK* | 1:1000 | *2532, Cell Signaling Technology* |
| *P-s588 TBC1D4* | 1:1000 | 8730, *Cell Signaling Technology* |
| *P-t642* | 1:1000 | 8881, *Cell Signaling Technology* |
| *TBC1D4* | 1:1000 | *(AS160), 07-741, Upstate Biotechnology, Inc, USA* |
| *P-s240/4 S6PK* | 1:1000 | *5364, Cell Signaling Technology* |
| *S6PK* | 1:1000 | *2217, Cell Signaling Technology* |
| *P-s65 4EBP1* | 1:1000 | *9456, Cell Signaling Technology* |
| *4EBP1* | 1:1000 | *9644, Cell Signaling Technology* |
| *Atrogin* | 1:1000 | *AP2041, ECM Biosciences, USA* |
| *Murf1* | 1:1000 | *ab172479, Abcam* |
| *LC3B* | 1:1000 | *L2543, Sigma Aldrich, Sweden* |
| *P-s256 FoxO1* | 1:1000 | *9461, Cell Signaling Technology* |
| *FoxO1* | 1:1000 | *2880, Cell Signaling Technology* |
| *P-s253 FoxO3a* | 1:1000 | *13129, Cell Signaling Technology* |
| *FoxO3a* | 1:1000 | *2497, Cell Signaling Technology* |
| *K48 ubiquitination* | 1:1000 | *Apu2; Millipore/ Sigma Aldrich, Sweden* |
| *Total ubiquitination* | 1:3000 | *Z0458; Dako, Sweden* |
| *Gadd45a* | 1:250 | *Sc-792; Santa Cruz Biotechnology* |
| *DDIT4* | 1:1000 | *10638-I-AP; Proteintech, UK* |
| *Rabbit-HRP* | 1:10-20000 | *7074, Cell Signaling Technology* |
| *Mouse-HRP* | 1:10-20000 | *Sc-2375, Santa Cruz Biotechnology* |
| *Rabbit-AF 594* | 1:1000 | *A11012, Invitrogen* |
| *Mouse- AF 488* | 1:1000 | A21121, Invitrogen |

**Supplementary Table 2.** Insulin Signaling gene transcripts following 8DES.

| Gene | Description | **logFC** (8DES vs 8D) | FDR |
|---|---|---|---|
| Calm2 | Calmodulin 2 | -1,025 | 0,006 |
| Eif4e2 | Eukaryotic translation initiation factor 4E family member 2 | -0,961 | 0,018 |
| Hk2 | Hexokinase 2 | -0,892 | 0,025 |
| G6pc3 | Glucose 6 phosphatase catalytic subunit 3 | -0,757 | 0,009 |
| Map2k1 | Mitogen activated protein kinase 1 | -0,633 | 0,012 |
| Ppp1cc | Protein phosphatase 1 catalytic subunit gamma | -0,463 | 0,011 |
| Flot2 | Flotillin 2 | 0,578 | 0,009 |
| Cblb | Cbl proto-oncogene B | 0,644 | 0,015 |
| Pik3r2 | Phosphoinositide-3-kinase regulatory subunit 2 | 0,726 | 0,020 |
| Inpp5a | Inositol polyphosphate-5-phosphatase A | 0,869 | 0,014 |
| Pygm | Glycogen phosphorylase, muscle associated | 1,010 | 0,007 |
| Prkag2 | Protein kinase AMP-activated non-catalytic subunit gamma 2 | 1,062 | 0,036 |
| Gys1 | Glycogen synthase 1 | 1,105 | 0,022 |
| Irs1 | Insulin receptor substrate 1 | 1,137 | 0,028 |
| Trip10 | Thyroid hormone receptor interactor 10 | 1,165 | 0,006 |
| Ppp1r3d | Protein phosphatase 1, regulatory subunit 3D | 1,246 | 0,015 |
| Flot1 | Flotillin 1 | 1,457 | 0,003 |
| Ppargc1a | PPARG coactivator 1 alpha | 1,481 | 0,049 |
| Fbp2 | Fructose-bisphosphatase 2 | 1,583 | 0,048 |
| Socs3 | Suppressor of cytokine signaling 3 | 2,460 | 0,043 |
| Tbc1d4 | TBC1 domain family, member 4 | 3,110 | 0,003 |

**Supplementary Table 3.** CamK gene transcripts promoted by 8DES.

| Gene | Description | **logFC** (8DES vs 8D) | FDR |
|---|---|---|---|
| Camk1d | calcium/calmodulin-dependent protein kinase ID | 1,465 | 0,047 |
| Camkk2 | calcium/calmodulin-dependent protein kinase 2 | 0,934 | 0,045 |

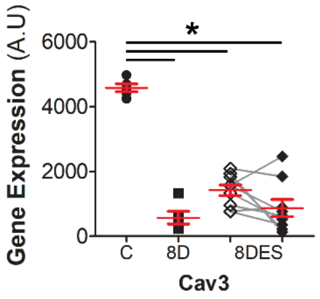


**Supplementary Figure 1. Cav3 RNA transcripts.** Adult Sprague Dawley rats were placed in critical care environment entailing deep sedation, CMV, and immobilization for 8 days with (8DES; Stimulated (STIM; open diamonds), Sham control (SHAM; closed diamonds); n = 9) and without (8D; closed squares; n =5) supramaximal ES and compared with control (CNT; filled circles; n = 5). Soleus RNA transcript expression of Cav3 (caveolin 3). *$p < 0.05$ significantly different from control.

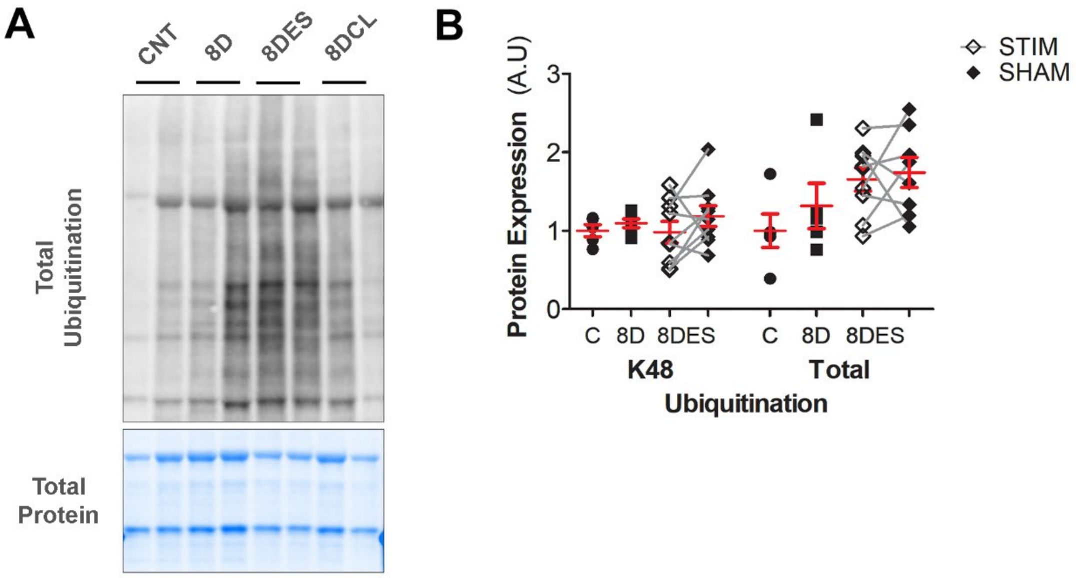


**Supplementary Figure 2 – Ubiquitination.** Adult female Sprague Dawley rats were placed in critical care environment entailing deep sedation, CMV, and immobilization for 8 days with (8DES; Stimulated (STM; open diamonds), Sham control (SHAM; closed diamonds); n = 9) and without (8D; closed squares; n = 5) supramaximal ES and compared with control (CNT; filled circles; n = 5). A) Representative westerns blotting of total Ubiquitination. B) Graphs showing densitometry measured in arbitrary units (A.U) of K48 and total ubiquitination.

**Methods**

*Animals and ICU Model*

All experimental animals were deeply sedated with isoflurane, pharmacologically paralyzed post-synaptically with α-cobratoxin, and mechanically ventilated at controlled volume as previously described (Fig 1A)(1). Adult female Sprague Dawley rats were divided into a control group (CNT; n=8), experimental group exposed to deep sedation, CMV, and neuromuscular blockade (NMB) for 8 days (8D; n=7), 8D with supramaximal soleus muscle ES (8DES; n=10). A subset of animals underwent 3 days of critical care conditions with (3DES; n = 3) or without (3D; n = 3) chronic supramaximal ES.

Isoflurane was delivered into the inspiratory gas stream by a precision mass-flow controller. During surgery, the anesthetic isoflurane was maintained > 1.5% Minimum Alveolar Concentration (MAC) to ensure: 1) high-voltage slow-wave activity electroencephalogram (EEG); 2) 90-100 mmHg mean arterial pressure and > 420 beats/min heart rate; and 3) no evident EEG, blood pressure, or heart rate responses to surgical manipulation. Post-operative Isoflurane was gradually lowered and maintained at <0.5% MAC. Rats were ventilated via a coaxial tracheal cannula at 72 breaths per min with an inspiratory and expiratory ratio of 1:2 and a minute volume of 180–200 ml. 40% $O_2$, 56,5% $N_2$, and 3% $CO_2$ gas concentrations of were delivered by a precision volumetric respirator. Airway pressure was monitored continuously to maintain normocapnic (end-tidal $CO_2$ = 37- 45 mmHg) and normoxic ($SpO_2$ > 90%) conditions, while also limiting stress and strain on the lungs. Intermittent hyperinflations (19-20 $cmH_2O$) over a constant positive end-expiratory pressure (1.5 cm $H_2O$) were set to prevent alveolar atelectasis. Post-synaptic neuromuscular blockade was induced on the first day (150 µg; α-cobratoxin) and maintained by continuous infusion (187 µg/day). CMV was initiated following NMB induction, avoiding hypercapnia and hypoxemia. Female rats were used to allow urine bladder catheterization via urethra and urination maintained >1 ml per h. In no case did animals show any signs of infection.

All experimental animals were maintained in fluid and nutritional balance throughout the experimental duration by introducing: 1) intra-arterial solution containing 21 ml $H_2O$, 24 ml 0.5 N lactated Ringers, 0.84 g oxacillin Na, 0.65 mg α-cobratoxin, 0.3 mg vitamin K (Synkavite), and 20 mg $K^+$ (as KCl); 2) an intra-venous solution (0.6 ml/h) containing 26 ml $H_2O$, 16 ml 0.5 N lactated Ringers, 20% glucose (Baxter, USA), and 0.32 g oxacillin Na for the initial 24 hours. After which 8.5% Travasol amino acids and 20% Intralipid (Kabi, Sweden) were added to the intra-venous solution to provide adequate nutrients (2, 3). Nutrient is provided by constant intra-venous delivery (0.6 ml/h). Body temperature, peripheral perfusion, and oxygen saturation were monitored by an infrared paw probe (MouseSTAT, Kent Scientific, USA) and maintained in physiological range. Female rats were used to allow urine bladder catheterization via urethra. The controls were anesthetized with isoflurane, maintained with spontaneous breathing, and sacrificed prior to tissue collection.

*Electrical stimulation*

While deeply sedated with isoflurane, hindlimbs of stimulated rats underwent surgical implantation of two Teflon-coated multistranded steel wires (outer diameter, 0.011 mm, AS 632, Cooner Sales, USA). The wires were placed across the soleus muscle, anterior and posterior of the motor point, as shown in

Figure 1A. (4-6). Both wires were fixed to the skin with sutures and connected to a constant current stimulator (701C; Aurora Scientific, Canada) and given supramaximal bipolar 0.2ms stimuli at constant current, typically between 10-30mA. Threshold for activation was monitored throughout the experiment, such that plantarflexion was maximal and any increase in stimulus did not increase ankle joint plantarflexion. The soleus muscle was stimulated for 10s every 20 s at 20Hz for 12h per day for 8 days and from this point on defined as 8DES Stimulated (STIM). The sham-operated contralateral limb, defined as SHAM 8DES, underwent electrode implantation, yet received no stimulation and controls for any potential surgical/implantation related effect.

*Single Muscle Fiber Contractile Measurements*

Following ICU conditions, deeply sedated rats were weighed and euthanized by heart removal. Soleus muscles were excised, weighed, and the mid belly region placed in relaxing solution at 4°C for muscle bundle preparation (4). The remaining muscle was snap frozen in liquid propane cooled by liquid nitrogen and stored at -140°C for further analyses. Muscle bundles were dissected free and tied with surgical silk to glass capillary tubes at 110% length. Bundles were then treated with skinning solution (relax solution containing glycerol) for 24 hours at 4°C before being transferred to -20°C. Following which, muscle bundles were subjected to increasing concentrations of sucrose solution (0.5-2 M), snap-frozen in liquid nitrogen-chilled propane, and maintained at -140°C. Prior to contractile assessment, bundles were subjected to decreasing concentrations of sucrose solution and kept in a skinning solution at −20°C. Single muscle fibers are carefully dissected from the muscle bundle by jewelers forceps and placed between two connectors. One connector leads to a force transducer (model 400A, Aurora Scientific, Canada) and the other to a lever arm system (model 308B, Aurora Scientific). The fiber were attached to the connectors as previously described (5). The apparatus was mounted on the stage of an inverted microscope (model IX70; Olympus, Sweden). While submerged in relaxing solution, the fiber sarcomere length was set to 2.65-2.75 µm. The diameter and depth of the fiber segment was measured at a magnification of ×320 with an image analysis system prior to the mechanical experiments. Fiber CSA was calculated from the diameter and depth, assuming an elliptical circumference, and was corrected for the 20% swelling known to occur during skinning (5).

For the mechanical recording, relaxing and activating solutions contained 4 mM Mg-ATP, 1 mM free $Mg^{2+}$, 20 mM imidazole, 7 mM EGTA, 14.5 mM creatine phosphate, and KCl to adjust the ionic strength to 180 mM and pH 7.0. The concentrations of free $Ca^{2+}$ were $10^{-9}$ M (relaxing solution) and $10^{-4.5}$ M (activating solution), expressed as $pCa^{2+}$ (-log [$Ca^{2+}$]). Apparent stability constants for $Ca^{2+}$-EGTA were corrected for temperature (15°C) and ionic strength (180 mM). The computer program from Fabiato was used to calculate the concentration of each metal, ligand, and metal-ligand complex (6). Immediately preceding each activation, the fiber was immersed for 15 s in a solution with a reduced $Ca^{2+}$-EGTA buffering capacity. This solution is identical to the relaxing solution except that the EGTA concentration is reduced to 0.5 mM, which results in a more rapid attainment of steady force during subsequent activation.

Force was calculated as the difference between the maximal steady-state isometric force in activating solution and the resting force measured in the same segment while in the relaxing solution. Maximal force production was normalized to CSA (specific force, absolute force ($P_0$)/CSA). A strict acceptance criterion was applied for contractile measurements. First, the sarcomere length was checked during the experiment, using a high-speed video analysis system (model 901A HVSL, Aurora Scientific). A muscle fiber was accepted and included in the analyses: (i) if the sarcomere length of a single muscle fiber changed <0.10 µm between relaxation and maximum activation, (ii) if maximal force changed <10% from first to seventh activation (5).

*Myosin: Actin Ratio*

Myosin and actin contents were quantified on a 12% SDS-PAGE gel. Electrophoresis was performed at 32 mA for 5 hours with a Tris-glycine electrode buffer (pH 8.3) at 15 °C (SE 600 vertical slab gel unit; Hoefer Scientific Instruments). Gels were stained using SimplyBlue SafeStain (Invitrogen, Sweden) and subsequently scanned in a soft laser densitometer (Molecular Dynamics, USA) with a high spatial resolution. The volume integration function was used to quantify the amount of protein and the results are presented as Myosin:Actin ratio (M:A Ratio).

*RNA Sequencing*

Soleus muscle was homogenized, and DNA disrupted by use of QIAshredder™ columns (Qiagen, Sweden), before RNA was extracted *via* Qiagen RNeasy® Fibrous tissue mini kit (Qiagen), as per the manufacturer's instructions. RNA quality and quantity were then quantified *via* 2100 Bioanalyzer (Agilent, Sweden) using an RNA 6000 Nano Kit (Agilent), as per the manufacturer's instructions. Strand-specific sequencing libraries were prepared from samples with an RNA Integrity Number (RIN) >8 using NEBNext® Ultra™ RNA Library Prep Kit for Illumina® (NEB, USA). Libraries were subject to sequencing on an Illumina platform (HiSeq Xten). After quality control, the paired-end clean reads were mapped to the rattus norvigicus Ensemble reference genome using HISAT2 (8). Differential gene expression and gene ontology analysis was undertaken using EdgeR and Goana pipeline on R (9). Genes with a false discovery rate (FDR) equal to $p \leq 0.05$ were reported as significantly differentially expressed genes (DEGs).

*Immunohistochemistry*

Eight-micrometer-thick frozen cross sections were cut from the mid-belly region of soleus muscles to assess GLUT4 translocation. Briefly, sections were fixed in acetone, permeabilized with 0.5% TBS-Tween and blocked with 1% BSA before reacted with an anti-GLUT4 and anti-dystrophin primary antibody in 1% BSA (Sup. Table 1). After which, samples were exposed to isotype specific anti-rabbit or mouse Alex Fluora fluorescent secondary antibodies and nuclei counterstained with DAPI. No primary antibody and no secondary antibody controls were included to confirm staining specificity. Representative images were captured by light microscopy at ×100 magnification.

*Periodic acid–Schiff (PAS) Immunohistochemistry*

Soleus cross sections were cut, fixed in Carnoy's fixative consisting of 60% ethanol, 30% chloroform and 10% Glacial acetic acid. Sections were washed and incubated in 0.5% Periodic Acid (Sigma) prior to exposure to Shiff's reagent (Sigma). Nuclei were counterstained with Mayer's hematoxylin (Sigma). Sections were then dehydrated and mounted. Representative images were captured by light microscopy (Zeiss) at ×400 magnification.

**References**


[Bibliography number]
